## Supplementary material for "Improved inference of population histories by integrating genomic and epigenomic data": SI Figures and Tables

### 1 Sequentially Markovian Coalescent for several markers

Our approach is a re-implementation of the PSMC',MSMC2, eSMC and eSMC2 but accounting for different genomic marker. Hence the Hidden Markov model is exactly the same as previously described, but with a different emission matrix. For each site, we first check what marker is present. We then set the correct substitution rate and number of hidden states. We note as *id* (identical) the event where both marker are in the same state, and *seg* if both are in different states (polymorphic). Extending the work in [1], we have the following formula:

$$
\begin{aligned}
P(id|\gamma) &= \frac{1}{nb_s} + \frac{(nb_s-1)}{nb_s} * e^{-2\mu t_\gamma \frac{(nb_s)}{(nb_s-1)}} \\
P(seg|\gamma) &= \frac{(nb_s-1)}{nb_s} - \frac{(nb_s-1)}{nb_s} * e^{-2\mu t_\gamma \frac{(nb_s)}{(nb_s-1)}}
\end{aligned}
\tag{1}
$$

Where $\mu$ is the substitution rate of the marker per N generation, $t\gamma$ the average coalescent time in state $\gamma$ and $nb_s$ the number of possible stat the marker can take.

### 2 Sequentially Markovian Coalescent with DNA methylation (SMCm)

SMCm is similar to PSMC',MSMC2, eSMC and eSMC2 but additionally accounting for epimutations.

#### 2.1 Accounting only for site DNA epimutations

Here we assume epimutations occur similarly as mutations (under a finite site model). Since the model accounts for sequence and DNA methylation polymorphisms, there are at each position 5 different possible observations when comparing two sequences. The first observation is 0 , corresponding to a non-methylable site where the two nucleotides are identical. 1, if the two nucleotide are different. 2 if it is a methylable site and both are unmethylated. 3, if the site is methylable and both are methylated. Finally, 4 is it is a methylable site and one cytosine is methylated and the other unmethylated. Therefore, after approximating the formula assuming DNA methylation state is not affected by DNA mutations we find:

$$
\begin{aligned}
P(0|\gamma) &= e^{-2\mu t_\gamma} \\
P(1|\gamma) &= 1 - e^{-2\mu t_\gamma} \\
P(2|\gamma) &= ((p_u \times (p_{m1} \times p_{m1})) + ((1-p_u) \times (1-p_{m2}) \times (1-p_{m2}))) \\
P(3|\gamma) &= ((p_u \times ((1-p_{m1}) \times (1-p_{m1}))) + ((1-p_u) \times (p_{m2}) \times (p_{m2}))) \\
P(4|\gamma) &= ((p_u \times (2 \times p_{m1} \times (1-p_{m1}))) + ((1-p_u) \times (2 \times p_{m2} \times (1-p_{m2})))) \\
p_u &= \frac{\mu_u}{\mu_u + \mu_m} \\
\theta_m &= (\mu_u + \mu_m) \times t_\gamma \\
p_{m1} &= (p_u + ((1-p_u) \times e^{(-\theta_m)})) \\
p_{m2} &= ((1-p_u) + (p_u \times e^{(-\theta_m)})))
\end{aligned} \tag{2}
$$

Where $\mu$ is the mutation rate per nucleotide per N generation,$\mu_m$ the methylation rate per generation,$\mu_u$ the demethylation rate per generation and $t_\gamma$ the average coalescent time in state $\gamma$. Additionally we define $p_u$ as the probability to be unmethylated at equilibrium, as well as $p_m1$ and $p_m2$ the respective probability to stay unmethelytaded or methylated after a time $t_\gamma$.

#### 2.2 Accounting only for region epimutations

Here we assume region epimutation occur similarly as mutations (under a finite site model). However, unlike previously, the epimutations affect multiple sites, hence only the first position of a methylated region is considered and the following positions will be considered as missing data (because it is one block of information). Therefore, there are at each position 6 different possible observations when comparing two sequences. The first observation is 0 , corresponding to a non-methylable site where the two nucleotides are identical. 1, if the two nucleotide are different. 2 if it is a region with methylation state annotated and both regions are unmethylated. 3,if it is a region with methylation state annotated and both regions are methylated. 4 if it is a region with methylation state annotated and both regions are in different methylation state. 5 is missing data. Therefore, after approximating the formula assuming methylation state is not affected by mutations we have the following formula:

$$
\begin{aligned}
P(0|\gamma) &= e^{-2\mu t_\gamma} \\
P(1|\gamma) &= 1 - e^{-2\mu t\gamma} \\
P(2|\gamma) &= ((p_u \times (p_{m1} \times p_{m1})) + ((1-p_u) \times (1-p_{m2}) \times (1-p_{m2}))) \\
P(3|\gamma) &= ((p_u \times ((1-p_{m1}) \times (1-p_{m1}))) + ((1-p_u) \times (p_{m2}) \times (p_{m2}))) \\
P(4|\gamma) &= ((p_u \times (2 \times p_{m1} \times (1-p_{m1}))) + ((1-p_u) \times (2 \times p_{m2} \times (1-p_{m2})))) \\
P(5|\gamma) &= 1 \\
p_u &= \frac{\mu_u}{\mu_u + \mu_m} \\
\theta_m &= (\mu_u + \mu_m) \times t_\gamma \\
p_{m1} &= (p_u + ((1-p_u) \times e^{(-\theta_m)})) \\
p_{m2} &= ((1-p_u) + (p_u * e^{(-\theta_m)}))) \qquad (3)
\end{aligned}
$$

Where $\mu$ is the mutation rate per nucleotide per N generation,$\mu_m$ the region methylation rate per generation,$\mu_u$ the region demethylation rate per generation and $t_\gamma$ the average coalescent time in state $\gamma$. Additionnally we define $p_u$ as the probability for the region to be unmethylated at equilibrium, as well as $p_m1$ and $p_m2$ the respective probability for the region to stay unmethelytaded or methylated after a time $t_\gamma$.

To recover the region epimutations we use a hidden markov model (HMM). The HMM takes as input genome and methylome data described as above, and compares two sequences to recover epiregion. The hidden markov model has 9 hidden states: 1 (regions with no methylation information), 2 (no methylation information in individual 1& mainly methylated region in individual 2) , 3 (no methylation information in individual 1& mainly unmethylated region in individual 2) , 4 (no methylation information in individual 2& mainly methylated region in individual 1) , 5 (mainly methylated region in individual 1& mainly methylated region in individual 2) , 6 (mainly methylated region in individual 1& mainly unmethylated region in individual 2) , 7 ( mainly unmethylated region in individual 1& no methylation information in individual 2) , 8 (mainly unmethylated region in individual 1& mainly methylated region in individual 2) , 9 (mainly unmethylated region in individual 1& mainly unmethylated region in individual 2). To define the transition rate, additional parameters are necessary. The user needs to define the minimum number of annotated methylable site to form a region (by default 4) and minimum size of a region in bp (100 by default). Our approach then defines the transition rate as the transition rate maximizing the number of regions respecting the defined criteria. The emission matrix is defined by the user, but by default the probability of observing methylated and unmethylated in a methylated region is respectively set to 0.8 and 0.2. In an unmethylated region, the probabilities are respectively set to 0.2 and 0.8. We use those values throughout the study.

#### 2.3 Accounting for site and region epimutations

Here we assume site and region epimutation occur similarly as mutations (under a finite site model). Like previously, the epimutations affect multiple sites, hence only the first position of a methylated region is considered and the following positions will be considered as missing data (because it is one block of information). However the noted observation will depend on the whole region. Therefore, there are at each position 10 different possible observations when comparing two sequences. The first observation is 0 , corresponding to a non-methylable site where the two nucleotides are identical. 1, if the two nucleotide are different. 2 if it is a region with methylation state annotated and both regions are unmethylated and all site are unmethylated. 3 if it is a region with methylation state annotated and both regions are unmethylated and all site are methylated. 4 if it is a region with methylation state annotated and both regions are unmethylated and at least one site is segregating. 5 if it is a region with methylation state annotated and both regions are methylated and all site are unmethylated. 6 if it is a region with methylation state annotated and both regions are methylated and all site are methylated. 7 if it is a region with methylation state annotated and both regions are methylated and at least one site is segregating. 8 if it is a region with methylation state annotated and both regions are in different methylation state. 9 is missing data. Therefore, we have the following formula:

$$
\begin{aligned}
P(0|\gamma) &= e^{-2\mu t_\gamma} \\
P(1|\gamma) &= 1 - e^{-2\mu t_\gamma} \\
P(2|\gamma) &= ((P_O(P_{OO}P_{OO})) + (P_P(P_{PO}P_{PO})) + (P_M(P_{MO}P_{MO})) + (P_U(P_{UO}P_{UO}))) \\
P(3|\gamma) &= ((P_O(P_{OP}P_{OP})) + (P_P(P_{PP}P_{PP})) + (P_M(P_{MP}P_{MP})) + (P_U(P_{UP}P_{UP}))) \\
P(4|\gamma) &= ((P_O(2P_{OP}P_{OO})) + (P_P(2P_{PP}P_{PO})) + (P_M(2P_{MP}P_{MO})) + (P_U(2P_{UP}P_{UO}))) \\
P(5|\gamma) &= ((P_O(P_{OU}P_{OU})) + (P_P(P_{PU}P_{PU})) + (P_M(P_{MU}P_{MU})) + (P_U(P_{UU}P_{UU}))) \\
P(6|\gamma) &= ((P_O(P_{OM}P_{OM})) + (P_P(P_{PM}P_{PM})) + (P_M(P_{MM}P_{MM})) + (P_U(P_{UM}P_{UM}))) \\
P(7|\gamma) &= ((P_O(2P_{OU}P_{OM})) + (P_P(2P_{PU}P_{PM})) + (P_M(2P_{MU}P_{MM})) + (P_U(2P_{UU}P_{UM}))) \\
P(8|\gamma) &= ((p_u(2p_{m1}(1-p_{m1}))) + ((1-p_u)(2p_{m2}(1-p_{m2})))) \\
P(9|\gamma) &= 1
\end{aligned}
\tag{4}
$$

Where we have :

$$
\begin{gathered}
p_u = \frac{\mu_u}{\mu_u + \mu_m}; p_{reg\_d} = \frac{\mu_{reg\_d}}{\mu_{reg\_d} + \mu_{reg\_m}} \\
\theta_m = (\mu_u + \mu_m)t_\gamma; \theta_{reg\_m} = (\mu_{reg\_d} + \mu_{reg\_m})t_\gamma \\
p_{m1} = (p_u + ((1 - p_u)e^{(-\theta_m)})); p_{m2} = ((1 - p_u) + (p_u e^{(-\theta_m)}))) \\
p_{reg\_m1} = (p_{reg\_d} + ((1 - p_{reg\_d})e^{(-\theta_{reg\_m})})); p_{reg\_m2} = ((1 - p_{reg\_d}) + (p_{reg\_d} e^{(-\theta_{reg\_m})}))) \\
t_m = \frac{1}{(\mu_{reg\_d} + \mu_{reg\_m})} \\
p_{eq1} = (p + ((1 - p)e^{(-(\mu_u + \mu_m)*(t_m))})); p_{eq2} = ((1 - p) + (pe^{(-(\mu_u + \mu_m)*(t_m))})) \\
P_{no\_reg\_event} = (exp(-(\mu_{reg\_d} + \mu_{reg\_m}) * t_\gamma)) \\
p_{m1\_small} = (p_u + ((1 - p_u)e^{(-(\mu_u + \mu_m)*((\frac{1}{(\mu_{reg\_d} + \mu_{reg\_m})}) - (\frac{(t_\gamma e^{(-(\mu_{reg\_d} + \mu_{reg\_m})t_\gamma)})}{(1 - exp(-(\mu_{reg\_d} + \mu_{reg\_m})t_\gamma))})))})) \\
p_{m2\_small} = ((1 - p_u) + (p_u e^{(-(\mu_u + \mu_m)*((\frac{1}{(\mu_{reg\_d} + \mu_{reg\_m})}) - (\frac{(t_\gamma e^{(-(\mu_{reg\_d} + \mu_{reg\_m})t_\gamma)})}{(1 - exp(-(\mu_{reg\_d} + \mu_{reg\_m})t_\gamma))})))})) \\
P_O = (p_{reg\_d})(1 - p_{eq1}) \\
P_P = (p_{reg\_d})(p_{eq1}) \\
P_U = (1 - p_{reg\_d})(1 - p_{eq2}) \\
P_M = (1 - p_{reg\_d})(p_{eq2}) \\
P_{OO} = p_{reg\_m1}(((1 - P_{no\_reg\_event})(1 - p_{m1\_small})) + (P_{no\_reg\_event}(p_{m2}))) \\
P_{OP} = p_{reg\_m1}(((1 - P_{no\_reg\_event})(p_M 1_s mall)) + (P_{no\_reg\_event}(1 - p_{m2}))) \\
P_{OM} = (1 - p_{reg\_m1})p_{m2\_small} \\
P_{OU} = (1 - p_{reg\_m1})(1 - p_{m2\_small}) \\
P_{PO} = p_{reg\_m1}(((1 - P_{no\_reg\_event})(1 - p_{m1\_small})) + (P_{no\_reg\_event}(1 - p_{m1}))) \\
P_{PP} = p_{reg\_m1}(((1 - P_{no\_reg\_event})(p_{m1\_small})) + (P_{no\_reg\_event}(p_{m1}))) \\
P_{PM} = (1 - p_{reg\_m1})p_{m2\_small} \\
P_{PU} = (1 - p_{reg\_m1})(1 - p_{m2\_small}) \\
P_{MO} = (1 - p_{reg\_m2})(1 - p_{m1\_small}) \\
P_{MP} = (1 - p_{reg\_m2})(p_{m1\_small}) \\
P_{MM} = p_{reg\_m2}(((1 - P_{no\_reg\_event})(p_{m2\_small})) + (P_{no\_reg\_event}(p_{m2}))) \\
P_{MU} = p_{reg\_m2}(((1 - P_{no\_reg\_event})(1 - p_{m2\_small})) + (P_{no\_reg\_event}(1 - p_{m2}))) \\
P_{UO} = (1 - p_{reg\_m2})(1 - p_{m1\_small}) \\
P_{UP} = (1 - p_{reg\_m2})(p_{m1\_small}) \\
P_{UM} = p_{reg\_m2}(((1 - P_{no\_reg\_event})(p_{m2\_small})) + (P_{no\_reg\_event}(1 - p_{m1}))) \\
P_{UU} = p_{reg\_m2}(((1 - P_{no\_reg\_event})(1 - p_{m2\_small})) + (P_{no\_reg\_event}(p_{m1})))
\end{gathered}
\tag{5}
$$

Where $\mu$ is the mutation rate per nucleotide per N generation,$\mu_m$ the site methylation rate per generation,$\mu_u$ the site demethylation rate,$\mu_{reg\_m}$ the region methylation rate per generation,$\mu_{reg\_d}$ the region demethylation rate per generation and $t_\gamma$ the average coalescent time in state $\gamma$. Please refer to the R function build_emi_m from the Package eSMC2 for a clearer description of the probabilities. Note that in the text, these probabilities are written as: $\mu_1$ for mutation rate per nucleotide, $\mu_{SM}$ the site methylation rate, $\mu_{SU}$ the site demethylation rate, $\mu_{RM}$ the region methylation rate, and $\mu_{RU}$ the region demethylation rate.

### 3 Supplementary Figures

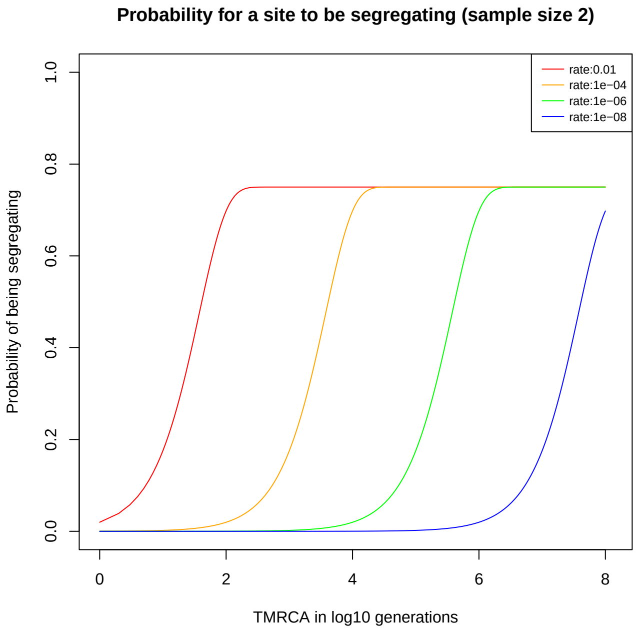


Supplementary Figure 1: **Probability of a site to be segregating in a sample of size two for different mutation rates**. The probability for a site to be segregating in a sample of size two under different mutation rates: $10^{-2}$ in red,$10^{-4}$ in orange, $10^{-6}$ in green and $10^{-8}$ in blue. The marker is assumed here to have $nb_s = 4$ possible states.

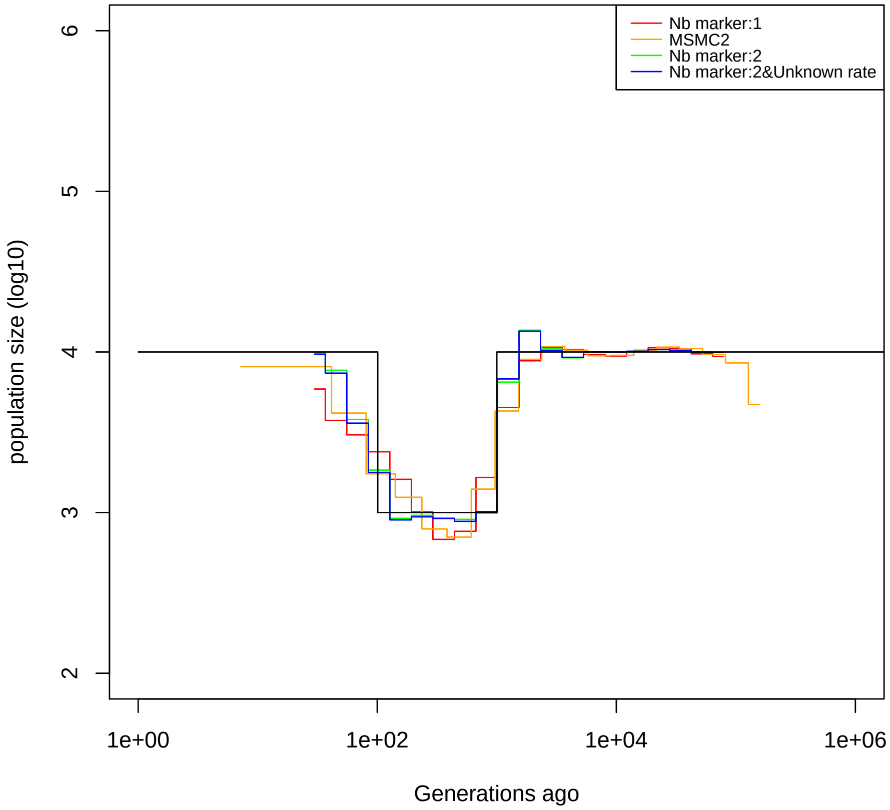


Supplementary Figure 2: **Performance of SMC using two theoretical markers and marker 2 is very rare.** Estimated demographic history of a recent bottleneck using theoretical genomic markers and 10 sequences of 100 Mb: with only marker 1 (red and orange) and with two markers (green and blue). Marker 2 is found at 0.1% of teh sites. The parameters are $r = 10^{-8}$ per generation per bp, and $\mu_1 = 10^{-8}$, $\mu_2 = 10^{-4}$ per generation per bp.

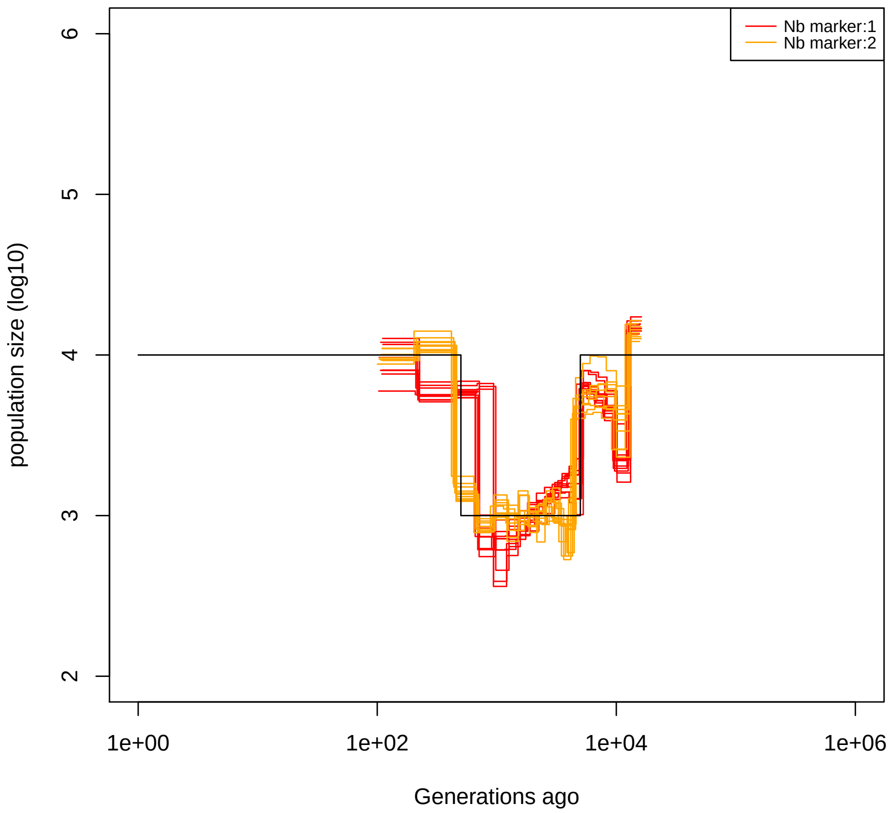


Supplementary Figure 3: **Performance of SMC using theoretical markers using the Likelihood estimation function of SMCtheo.** Estimated demographic history by SMC on theoretical genomic markers using 6 scaffolds each of 20 Mb with sample size 10: with one marker in red, and two markers in orange. We use here the likelihood function (LH) estimation procedure from SMCtheo. The parameters are $r = 10^{-8}$ per generation per bp, and $\mu_1 = 10^{-8}$, $\mu_2 = 10^{-4}$ per generation per bp.

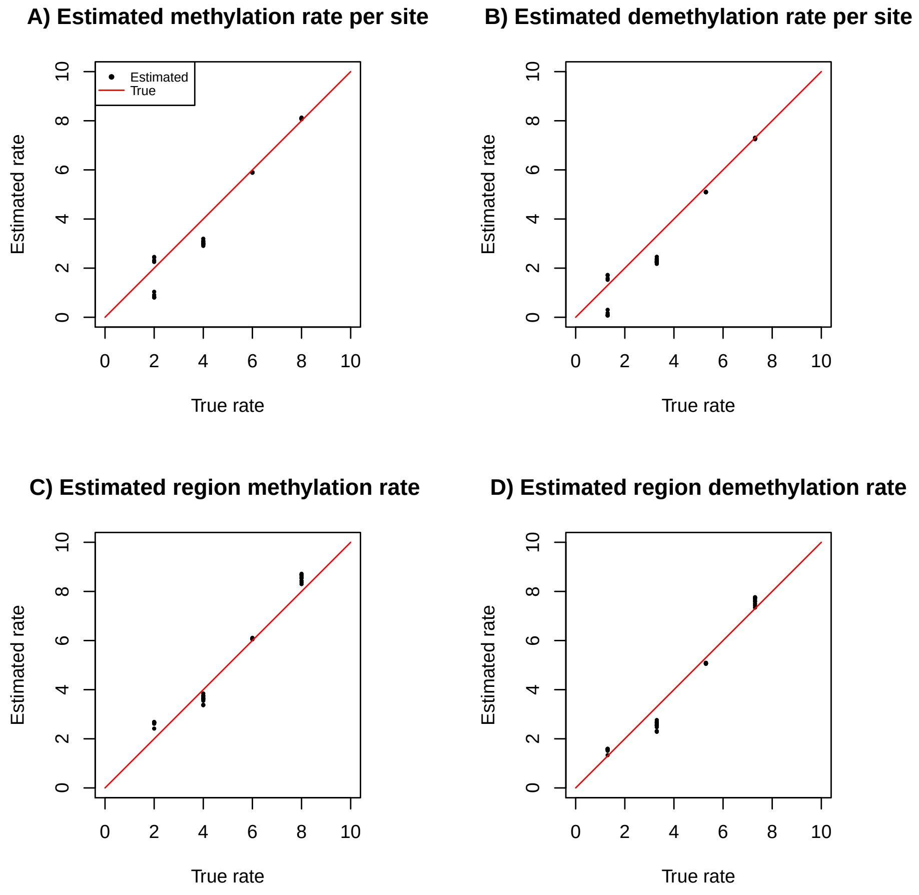


Supplementary Figure 4: **Average estimates of the site and region methylation and demethylation rates for simulated data.** The true rate is indicated as x-axis and the estimated in y-axis in log10 scale. We use ten repetitions with 10 sequences of 100 Mb with $r = \mu_1 = 10^{-8}$ per generation per bp under a constant population size fixed to 10,000.

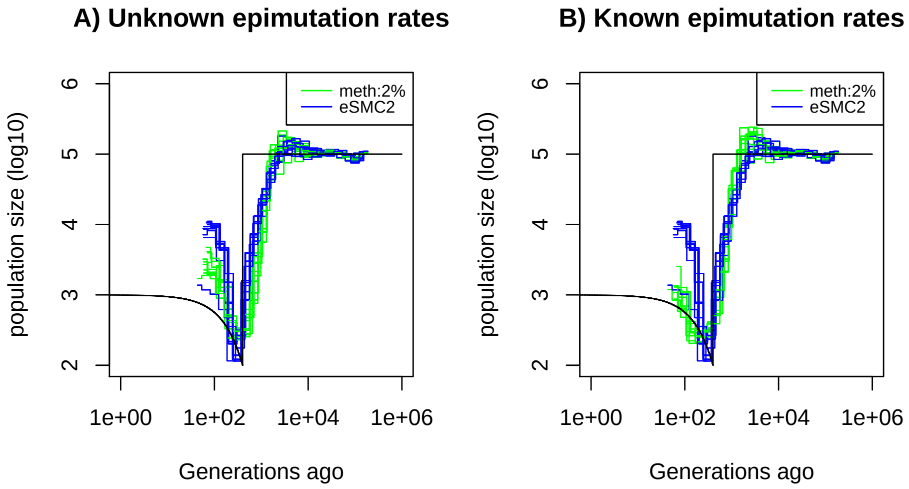


Supplementary Figure 5: **Performance of SMCm for methylation with only DMR regions of length 1kbp.** Estimated demographic history by eSMC2 (blue) and SMCm in presence of region epimutations only and of length 1kbp (green) using 10 sequences of 100 Mb under a recent bottleneck (black). The parameters are $r = 3.5 \times 10^{-8}$ per generation per bp, $\mu_1 = 7 \times 10^{-9}$, and the region methylation $\mu_{RM} = 2 \times 10^{-4}$ and demethylation rate $\mu_{RU} = 1 \times 10^{-3}$ per generation per bp.

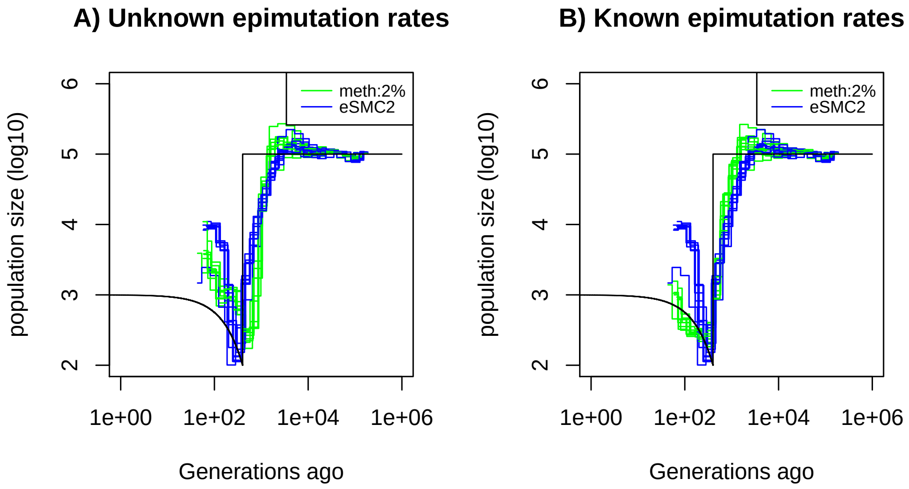


Supplementary Figure 6: **Performance of SMCm for methylation with only DMR regions of length 150bp** Estimated demographic history by eSMC2 (blue) and SMCm in presence of region epimutations only and of length 150bp (green) using 10 sequences of 100 Mb under a recent bottleneck (black). The parameters are $r = 3.5 \times 10^{-8}$ per generation per bp, $\mu_1 = 7 \times 10^{-9}$, and the region methylation $\mu_{RM} = 2 \times 10^{-4}$ and demethylation rate $\mu_{RU} = 1 \times 10^{-3}$ per generation per bp.

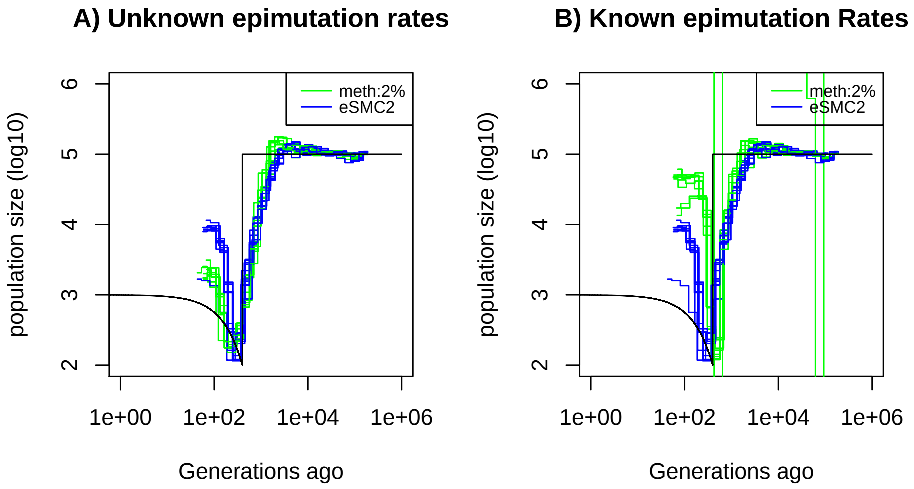


Supplementary Figure 7: **Performance of SMCm for methylation with site and region epimutations.** Estimated demographic history by eSMC2 (blue) and SMCm in presence of site and region epimutations (green) using 10 sequences of 100 Mb under a recent bottleneck (black). The recombination is set to $r = 3.5 \times 10^{-8}$ per generation per bp, the mutation rate is set to $\mu_1 = 7 \times 10^{-9}$, site methylation rate to $\mu_{SM} = 3.5 \times 10^{-4}$, site demethylation rate to $\mu_{SU} = 1.5 \times 10^{-3}$ per generation per bp, region methylation rate to $\mu_{RM} = 2 \times 10^{-4}$ and region demethylation rate to $\mu_{RU} = 1 \times 10^{-3}$ per generation per bp.

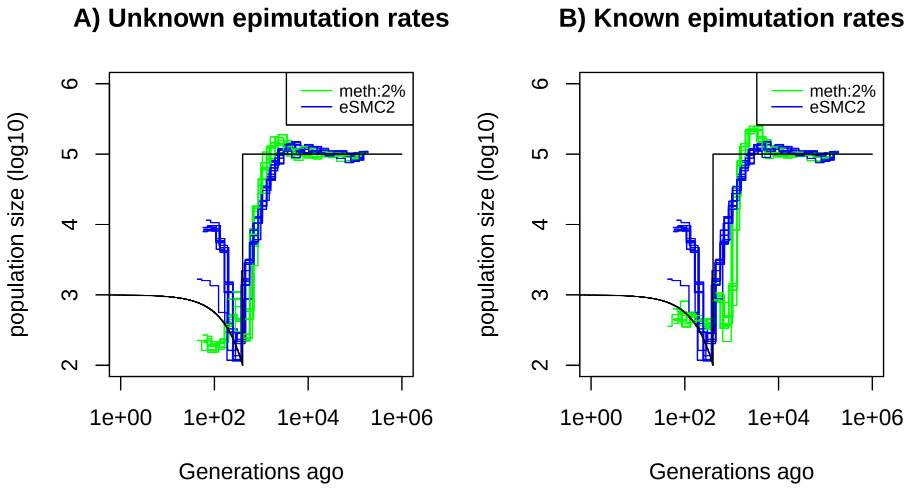


Supplementary Figure 8: **Performance of SMCm for methylation, accounting only for SMPs.** Estimated demographic history by eSMC2 (blue) and SMCm in presence of site and region epimutations but only accounting for site epimutations SMPs (green) using 10 sequences of 100 Mb under a recent bottleneck (black). The recombination is set to $r = 3.5 \times 10^{-8}$ per generation per bp, the mutation rate is set to $\mu_1 = 7 \times 10^{-9}$, site methylation rate to $\mu_{SM} = 3.5 \times 10^{-4}$, site demethylation rate to $\mu_{SU} = 1.5 \times 10^{-3}$ per generation per bp, region methylation rate to $\mu_{RM} = 2 \times 10^{-4}$ and region demethylation rate to $\mu_{RU} = 1 \times 10^{-3}$ per generation per bp.

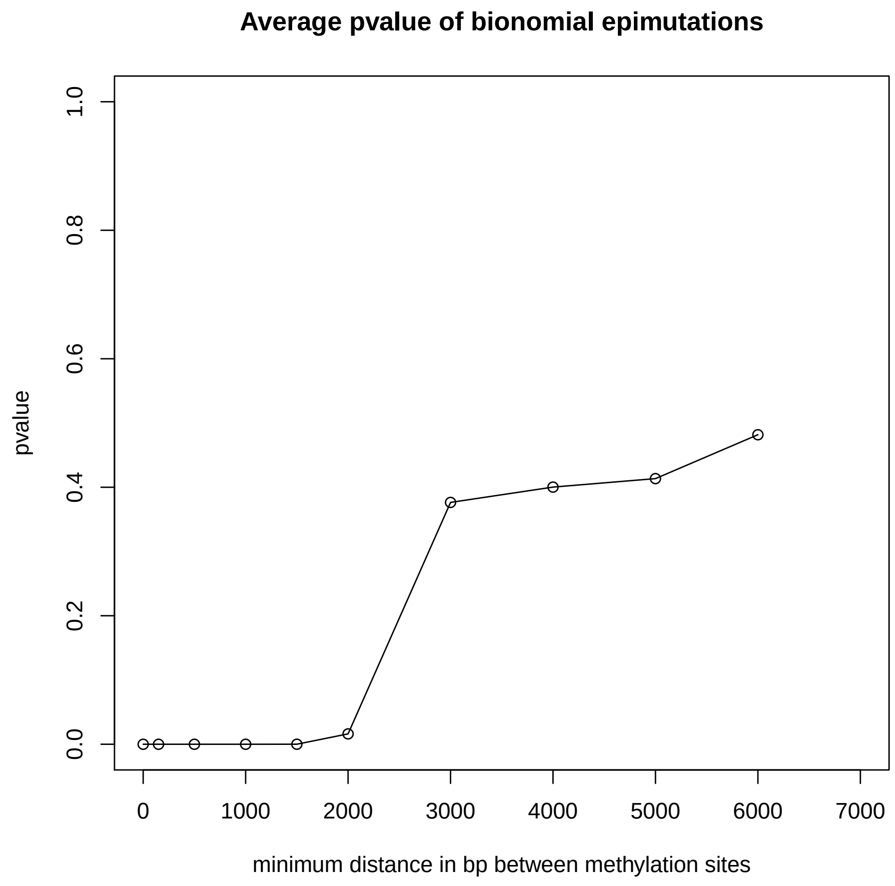


Supplementary Figure 9: **Average pvalue of the binomial test for epimutations.** Average p-value across 10 genomes of the binomial test on epimutations seperated by a minimum distance in bp (x axis) on our eight methylome scaffolds of *A. thaliana*

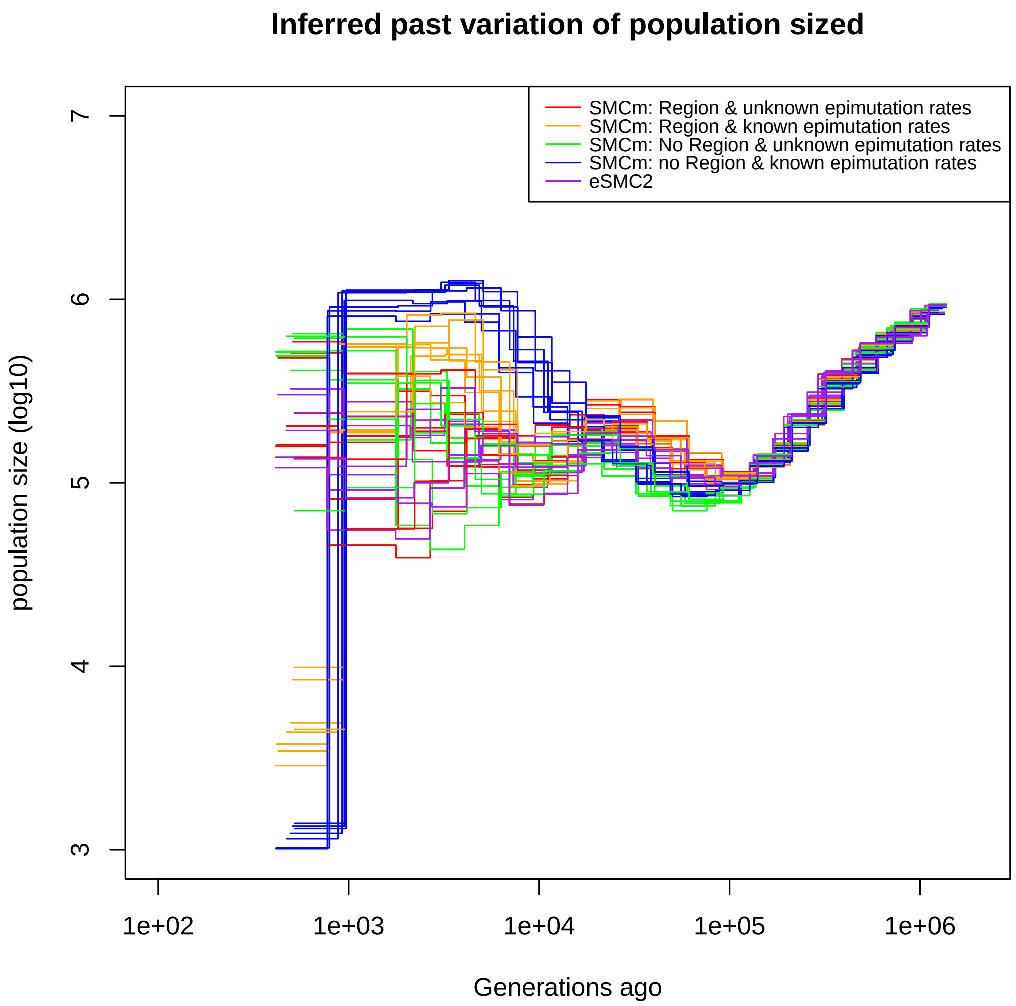


Supplementary Figure 10: **Demographic estimation using all methylation sites from German accessions of *A. thaliana*.** Estimated demographic history of the German population by eSMC2 (purple) and SMCm under different assumptions. In green the SMCm results with epimutation rates estimated and regions epimutation (DMRs) ignored. In blue are the estimates with epimutation rates fixed and regions epimutation (DMRs) ignored. In red are the results with epimutation rates sites and regions estimated by SMCm, and in orange with both regions and site epimutation fixed to empirical values. The recombination is set to $r = 3.6 \times 10^{-8}$ per generation per bp, the mutation rate is set to $\mu_1 = 6.95 \times 10^{-9}$, site methylation rate to $\mu_{SM} = 3.5 \times 10^{-4}$ and site demethylation rate to $\mu_{SU} = 1.5 \times 10^{-3}$ per generation per bp (when fixed). When fixed, the region methylation and demethylation rates are set, respectively, to $\mu_{RM} = 1.6 \times 10^{-4}$ and $\mu_{RU} = 9.5 \times 10^{-4}$.

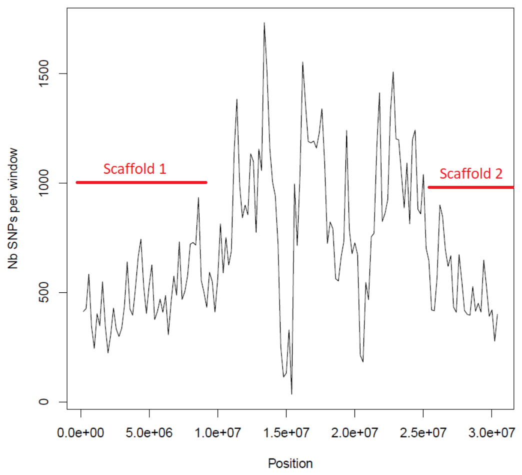


Supplementary Figure 11: **Average number of segregating site per window of 100kp on chromosome 1** Estimated Average number of segregating site per window of 100kp on chromosome 1 on 10 individuals of the German accessions (black).

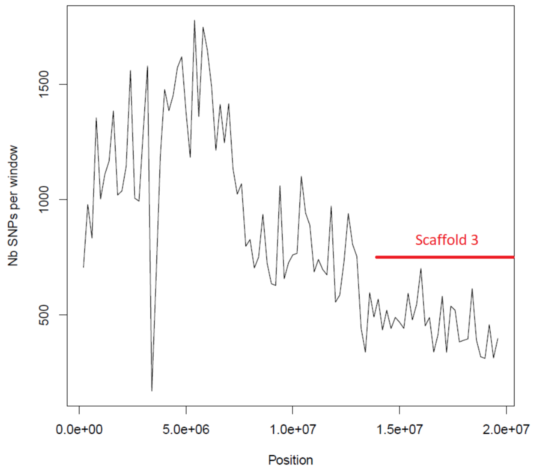


Supplementary Figure 12: **Average number of segregating site per window of 100kp on chromosome 2** Estimated Average number of segregating site per window of 100kp on chromosome 2 on 10 individuals of the German accessions (black).

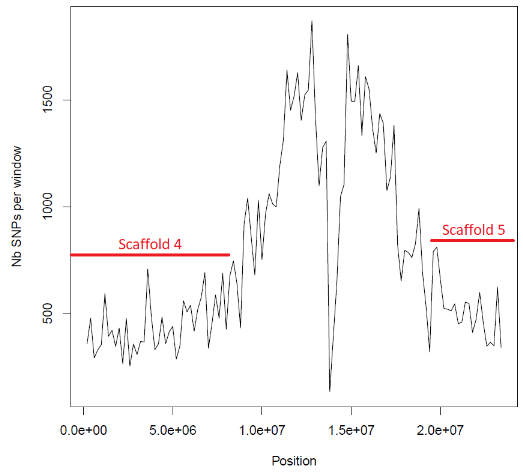


Supplementary Figure 13: **Average number of segregating site per window of 100kp on chromosome 3** Estimated Average number of segregating site per window of 100kp on chromosome 3 on 10 individuals of the German accessions (black).

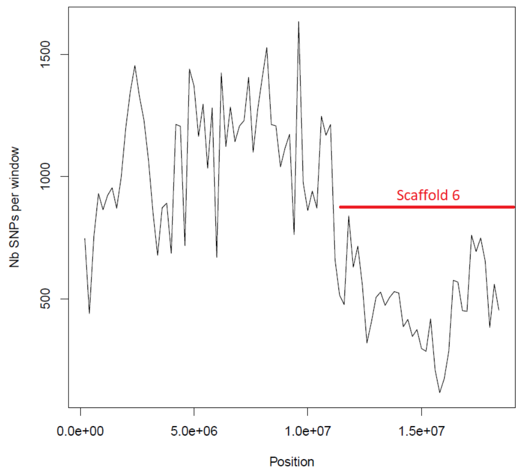


Supplementary Figure 14: **Average number of segregating site per window of 100kp on chromosome 4** Estimated Average number of segregating site per window of 100kp on chromosome 4 on 10 individuals of the German accessions (black).

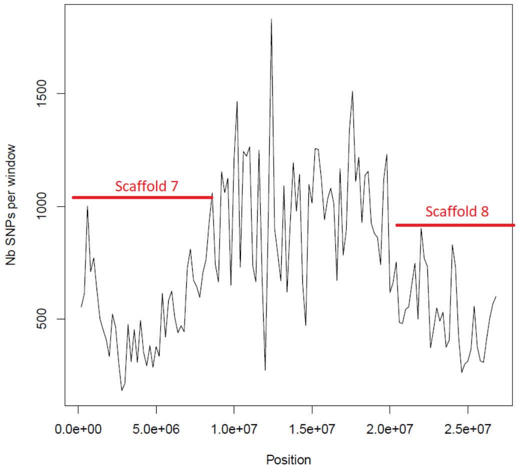


Supplementary Figure 15: **Average number of segregating site per window of 100kp on chromosome 5** Estimated Average number of segregating site per window of 100kp on chromosome 5 on 10 individuals of the German accessions (black).

### 4 Supplementary Tables

| Approach | MRSE Fig 2A | MRSE Fig 2B | MRSE Fig 2C | MRSE Fig 2D | MRSE Sup. Fig 2 | MRSE Sup. Fig 3 |
|---|---|---|---|---|---|---|
| eSMC2 | 6.92 | 5.41 | 8.68 (0.020) | 9.17 | 6.92 | 7.12 (0.01) |
| SMCtheo (known rates) | 6.42 | 6.00 | 6.77 (0.004) | 8.74 | 6.73 | 6.90 (0.02) |
| SMCtheo (unknown rates) | 6.60 | 5.98 | | 10.59 | 6.73 | |
| MSMC2 | 7.02 | 10.1 | 11.76 (0.04) | 10.74 | 6.83 | |

Supplementary Table 1: Average mean root square error (in log10) of demographic inference in Figure 2A-D, Sup. Figure 2 and Sup. Figure 3 by the three approach (eSMC2, SMCtheo with unknown rates, SMCtheo with known rates and MSMC2). The coefficient of variation is indicated in parentheses

| Region methylation rate | Region demethylation rate | Site methylation rate | Site demethylation rate | % of repetions accepting region epimutations |
|---|---|---|---|---|
| $1 \times 10^{-2}$ | $5 \times 10^{-2}$ | 0 | 0 | 100% |
| $1 \times 10^{-4}$ | $5 \times 10^{-4}$ | 0 | 0 | 100% |
| $1 \times 10^{-6}$ | $5 \times 10^{-6}$ | 0 | 0 | 100% |
| $1 \times 10^{-8}$ | $5 \times 10^{-8}$ | 0 | 0 | 100% |
| 0 | 0 | $1 \times 10^{-8}$ | $5 \times 10^{-8}$ | 2% |
| 0 | 0 | $1 \times 10^{-6}$ | $5 \times 10^{-6}$ | 1% |
| 0 | 0 | $1 \times 10^{-4}$ | $5 \times 10^{-4}$ | 0% |
| 0 | 0 | $1 \times 10^{-2}$ | $5 \times 10^{-2}$ | 0% |
| $1 \times 10^{-4}$ | $5 \times 10^{-4}$ | $1 \times 10^{-8}$ | $5 \times 10^{-8}$ | 100% |
| $1 \times 10^{-4}$ | $5 \times 10^{-4}$ | $1 \times 10^{-6}$ | $5 \times 10^{-6}$ | 100% |
| $1 \times 10^{-4}$ | $5 \times 10^{-4}$ | $1 \times 10^{-4}$ | $5 \times 10^{-4}$ | 100% |
| $1 \times 10^{-4}$ | $5 \times 10^{-4}$ | $1 \times 10^{-2}$ | $5 \times 10^{-2}$ | 0% |
| $1 \times 10^{-4}$ | $5 \times 10^{-4}$ | $5 \times 10^{-8}$ | $1 \times 10^{-8}$ | 100% |
| $1 \times 10^{-4}$ | $5 \times 10^{-4}$ | $5 \times 10^{-6}$ | $1 \times 10^{-6}$ | 100% |
| $1 \times 10^{-4}$ | $5 \times 10^{-4}$ | $5 \times 10^{-4}$ | $1 \times 10^{-4}$ | 100% |
| $1 \times 10^{-4}$ | $5 \times 10^{-4}$ | $5 \times 10^{-2}$ | $1 \times 10^{-2}$ | 0% |

Supplementary Table 2: Percentage of repetitions rejecting the $H_0$ hypothesis at p=0.05 of binomial distribution of epimutations over 100 repetitions using two sequences of 100 Mb with recombination and mutation rate set to $1 \times 10^{-8}$ per generation per bp under a constant population size fixed to 10,000.

| True site methylation rate | Estimated site methylation rate | True site demethylation rate | Estimated site demethylation rate |
|---|---|---|---|
| $10^{-8}$ | $1.0 \times 10^{-8}$ (0.03) | $5 \times 10^{-8}$ | $5.0 \times 10^{-8}$ (0.03) |
| $10^{-6}$ | $1.10^{-6}$ (0.01) | $5 \times 10^{-6}$ | $5.3 \times 10^{-6}$ (0.01) |
| $10^{-4}$ | $1.6 \times 10^{-4}$ (0.05) | $5 \times 10^{-4}$ | $8 \times 10^{-4}$ (0.05) |
| $10^{-2}$ | $3.6 \times 10^{-3}$ (0.74) | $5 \times 10^{-2}$ | $1.8 \times 10^{-2}$ (0.74) |

Supplementary Table 3: True versus average estimated values of the site methylation and demethylation rates over ten repetitions. We use two sequences of 100 Mb with $r = \mu_1 = 10^{-8}$ per generation per bp under a constant population size fixed to 10,000. The coefficient of variation is indicated in brackets.

| True region methylation rate | Estimated region methylation rate | True region demethylation rate | Estimated region demethylation rate |
|---|---|---|---|
| $10^{-8}$ | $6.3 \times 10^{-9}$ (0.02) | $5 \times 10^{-8}$ | $3.10^{-8}$ (0.02) |
| $10^{-6}$ | $1.10^{-6}$ (0.03) | $5 \times 10^{-6}$ | $5.4 \times 10^{-6}$ (0.03) |
| $10^{-4}$ | $1.6 \times 10^{-4}$ (0.16) | $5 \times 10^{-4}$ | $7.8 \times 10^{-4}$ (0.15) |
| $10^{-2}$ | $2.6 \times 10^{-3}$ (0.77) | $5 \times 10^{-2}$ | $1.4 \times 10^{-2}$ (0.77) |

Supplementary Table 4: True versus average estimated values of the region methylation and demethylation rates over ten repetitions. We use two sequences of 100 Mb with $r = \mu_1 = 10^{-8}$ per generation per bp under a constant population size fixed to 10,000. The coefficient of variation is indicated in brackets

| Methylation rate | Estimated methylation rate | Demethylation rate | Estimated demethylation rate |
|---|---|---|---|
| $10^{-8}$ | $8.0 \times 10^{-9}$ (0.04) | $5 \times 10^{-8}$ | $5.2 \times 10^{-8}$ (0.04) |
| $10^{-6}$ | $1.3 \times 10^{-6}$ (0.01) | $5 \times 10^{-6}$ | $8.0 \times 10^{-6}$ (0.01) |
| $10^{-4}$ | $9.5 \times 10^{-4}$ (0.19) | $5 \times 10^{-4}$ | $5.1 \times 10^{-3}$ (0.19) |
| $10^{-2}$ | $8.5 \times 10^{-2}$ (0.85) | $5 \times 10^{-2}$ | $4.6 \times 10^{-1}$ (0.85) |
| Region methylation rate | Estimated region methylation rate | Region demethylation rate | Estimated region demethylation rate |
| $10^{-8}$ | $2.9 \times 10^{-9}$ (0.37) | $5 \times 10^{-8}$ | $2.6 \times 10^{-8}$ (0.37) |
| $10^{-6}$ | $8.4 \times 10^{-7}$ (0.03) | $5 \times 10^{-6}$ | $8.4 \times 10^{-6}$ (0.03) |
| $10^{-4}$ | $2.5 \times 10^{-4}$ (0.38) | $5 \times 10^{-4}$ | $3.0 \times 10^{-3}$ (0.38) |
| $10^{-2}$ | $2.3 \times 10^{-3}$ (0.22) | $5 \times 10^{-2}$ | $2.9 \times 10^{-2}$ (0.22) |

Supplementary Table 5: Average estimated values of the site and region methylation and demethylation rates over ten repetitions using 2 sequences of 100 Mb with recombination and mutation rate set to $1 \times 10^{-8}$ per generation per bp under a constant population size fixed to 10,000. The coefficient of variation is indicated in brackets.

| Approach | MRSE Fig 5A | MRSE Fig 5B | MRSE Fig 5C | MRSE Fig 5D |
|---|---|---|---|---|
| eSMC2 | 9.18 (0.004) | 9.18 (0.004) | 9.44 (0.034) | 9.44 (0.034) |
| SMCm (2%) | 9.34 (0.013) | 9.22 (0.037) | 9.79 (0.057) | 9.59 (0.034) |
| SMCm (10%) | 9.31 (0.07) | 9.12 (0.035) | () | () |
| SMCm (20%) | 9.26(0.011) | 9.07 (0.023) | () | () |
| Approach | MRSE Fig 5A (gen<400) | MRSE Fig 5B (gen<400) | MRSE Fig 5C (gen<400) | MRSE Fig 5D (gen<400) |
| eSMC2 | 7.12 (0.098) | 7.12 (0.098) | 7.01 (0.12) | 7.01 (0.12) |
| SMCm (2%) | 6.46 (0.040) | 4.27 (0.13) | 6.12 (0.17) | 5.00 (0.12) |
| SMCm (10%) | 6.41 (0.045) | 4.43 (0.14) | () | () |
| SMCm (20%) | 6.51 (0.043) | 4.48 (0.10) | () | () |

Supplementary Table 6: Average mean root square error (in log10) of demographic inference in Figure 5 by the two approaches eSMC2, SMCm with unknown epimutations rates (A and C), and SMCm with known epimutation rates (B and D). Note the Second row indicates the MRSE in recent times (younger than 400 generations ago). The coefficient of variation is indicated in parentheses

| Approach | Mean Root Square Error |
|---|---|
| eSMC2 | 46,354 (0.20) |
| SMCm (known epimutation rates) | 46,317 (0.20) |
| SMCm (unknown epimutation rates) | 49,433 (0.22) |

Supplementary Table 7: Average mean root square error of inferred coalescent time (in generation unit) along the genome over ten repetitions by the three approaches (eSMC2, SMCm with unknown epimutation rates and SMCm with known epimutation rates) under the same scenario from Figure 5. Inference was performed on two haploid sequences of 10 Mb with $\mu = 7 \times 10^{-9}$, $r = 3.5 \times 10^{-8}$ per generation per bp. Methylation and demethylation rates were respectively fixed to $3.5 \times 10^{-4}$ and $1.5 \times 10^{-3}$ per generation per bp. The selfing rate was fixed to 90%. The coefficient of variation is indicated in parentheses

| Approach | MRSE Sup Fig 5A | MRSE Sup Fig 5B | MRSE Sup Fig 6A | MRSE Sup Fig 6B |
|---|---|---|---|---|
| eSMC2 | 9.21 (0.006) | 9.21 (0.006) | 9.22 (0.010) | 9.22 (0.010) |
| SMCm (2%) | 9.26 (0.007) | 9.30 (0.014) | 9.27 (0.017) | 9.59 (0.034) |
| Approach | MRSE Sup Fig 5A (gen<400) | MRSE Sup Fig 5B (gen<400) | MRSE Sup Fig 5C (gen<400) | MRSE Sup Fig 5D (gen<400) |
| eSMC2 | 7.21 (0.10) | 7.21 (0.10) | 7.34 (0.065) | 7.34 (0.065) |
| SMCm (2%) | 6.21 (0.046) | 4.87 (0.008) | 6.41 (0.048) | 4,72 (0.064) |

Supplementary Table 8: Average mean root square error (in log10) of demographic inference in Sup Figure 5 and 6 by the three approaches (eSMC2, SMCm with unknown epimutations rates and SMCm with known epimutation rates). Note the Second row indicates the MRSE in recent times (younger than 400 generations ago). The coefficient of variation is indicated in parentheses

| Approach | MRSE Sup Fig 7A | MRSE Sup Fig 7B |
|---|---|---|
| eSMC2 | 9.19 (0.002) | 9.19 (0.002) |
| SMCm (2%) | 9.19 (0.005) | 9.75 (0.16) |
| Approach | MRSE Fig 7A (gen<400) | MRSE Fig 7B (gen<400) |
| eSMC2 | 7.27 (0.08) | 7.27 (0.08) |
| SMCm (2%) | 5.88 (0.03) | 9.61 (0.18) |

Supplementary Table 9: Average mean root square error (in log10) of demographic inference in Sup Figure 7 by the three approaches (eSMC2, SMCm with unknown epimutations rates, SMCm with known epimutation rates). Note the Second row indicates the MRSE in recent times (younger than 400 generations ago). The coefficient of variation is indicated in parentheses

| Approach | MRSE Sup Fig 8A | MRSE Sup Fig 8B |
|---|---|---|
| eSMC2 | 9.19 (0.002) | 9.19 (0.002) |
| SMCm (2%) | 9.15 (0.007) | 9.47 (0.007) |
| Approach | MRSE Fig 8A (gen<400) | MRSE Fig 8B (gen<400) |
| eSMC2 | 7.27 (0.08) | 7.27 (0.08) |
| SMCm (2%) | 5.15 (0.04) | 4.60 (0.02) |

Supplementary Table 10: Average mean root square error (in log10) of demographic inference in Sup Figure 8 by the three approaches (eSMC2, SMCm with unknown epimutations rates, SMCm with known epimutation rates). Note the Second row indicates the MRSE in recent times (younger than 400 generations ago). The coefficient of variation is indicated in parentheses

| Methylation sites chosen | Estimated site methylation rate | Estimated site demethylation rate |
|---|---|---|
| polymorphic SMPs CG sites | $4.4 \times 10^{-5}$ | $5.9 \times 10^{-5}$ |
| CG sites separated by 3,000bp | $2.5 \times 10^{-7}$ | $2.1 \times 10^{-6}$ |
| all annotated CG sites | $4.4 \times 10^{-7}$ | $2.1 \times 10^{-6}$ |

Supplementary Table 11: Average estimated values of the site methylation and demethylation rates by SMCm using genomes and methylomes from 10 German accessions of *A. thaliana*. We use eight scaffolds each of 10 sequences with recombination and mutation rate respectively set to $r = 3.6 \times 10^{-8}$ and $\mu_1 = 6.95 \times 10^{-9}$ per generation per bp with selfing set to 90%. The polymorphic SMPs CG sites estimations corresponds to the green line in Figure 6. All CG sites estimations and CG site separated by 3,000bp corresponds to the data of the green line in Supplementary Figure 10.

| Methylation | Estimated methylation rate | Estimated demethylation rate |
|---|---|---|
| sites | $9 \times 10^{-8}$ | $4.9 \times 10^{-7}$ |
| regions | $2.0 \times 10^{-7}$ | $7.3 \times 10^{-7}$ |

Supplementary Table 12: Average estimated values of the site and region methylation and demethylation rates by SMCm using genomes and methylomes from 10 German accessions of *A. thaliana*. These estimations are produced during the inference of the red line in Supplementary Figure 10. We use eight scaffolds each of 10 sequences with recombination and mutation rate respectively set to $r = 3.6 \times 10^{-8}$ and $\mu_1 = 6.95 \times 10^{-9}$ per generation per bp with selfing set to 90%.
